# Synchrotron phase contrast micro-CT of prostate tissue

**DOI:** 10.64898/2026.08.12.742892

**Authors:** Roger M. Bourne, Benedicta Arhatari, Geoffrey Watson, Timur Gureyev, Adam Phipps, Samson Dowland, Nyoman D. Kurniawan, Paul Sved

## Abstract

Formalin-fixed prostate tissue samples were imaged by propagation-based synchrotron phase contrast micro computed tomography (*µ*CT) with a 3D spatial resolution of ca. 3 *µ*m. Post-*µ*CT, samples were prepared for histology with sections close to coplanar with the transverse *µ*CT image planes. Haematoxylin and eosin stained sections were examined by an expert prostate histopathologist and compared qualitatively with corresponding *µ*CT-visible microstructure features. There is potential for *µ*CT to provide complimentary information to conventional histology and light microscopy without the need for preparation of stained thin sections. For the imaging conditions and spatial resolution of our study, *µ*CT may provide tissue architectural features similar to those used in Gleason grading, albeit without clear subcellular microstructure detail. At the spatial resolution of our study *µ*CT may provide novel 3D microstructure information for validation of diffusion weighted magnetic resonance imaging (MRI) methods. As an example, we demonstrate a qualitative correlation between *µ*CT-derived stromal fibre orientation and preferential water diffusion direction measured by diffusion tensor MRI microscopy of the same sample.

## Introduction

Phase contrast micro computed tomography (*µ*CT) is gaining increasing interest for its ability to provide 3D cellular or subcellular soft tissue structural information on a par with the inherently 2D processes of traditional histology and light microscopy of thin sections. Reconstruction of 3D volumes from thin section histological techniques destroys the tissue sample, introduces additional artifacts, and requires tedious and time-consuming processing of large numbers of serial sections. Since all biological structures and processes occur in a 3D space, there are fundamental advantages in non-destructive 3D tissue characterisation methods such as *µ*CT.

Notwithstanding the advantages of *µ*CT, its contrast mechanism is physically different from those used in the histological processes that form the basis of traditional tissue characterisation and disease definitions. The current requirement for synchrotron light sources for phase contrast *µ*CT precludes applications in routine clinical specimen analysis but nevertheless provides opportunities for basic science applications, and for creation of an evidence base for development of in vivo clinical imaging applications such as breast cancer assessment^1^. Although lower cost “benchtop” light sources are available, these systems are currently limited by relatively low light intensity and coherence, with consequent long imaging times and limited soft tissue contrast^2^. The spatial coherence of synchrotron radiation enables phase-contrast imaging with high sensitivity to small density variations in low-density materials such as soft tissue^3^, without addition of X-ray dense contrast enhancement agents. Moreover, the high brightness of a synchrotron light source enables *µ*CT imaging to be performed in a very short time. Histology serves as the gold standard for cancer diagnosis. However, there is a need to perform direct comparisons of *µ*CT and histology in order to determine what histological features are available from *µ*CT, and whether *µ*CT can provide additional structural information not available from histology. Such *µ*CT-histology comparisons, often referred to as ‘virtual histology’, have been performed in diverse tissues including breast^4,5^, pancreas^6^, brain^7^, lung^8^, oesophagus^9^, kidney^10^, and prostate^10–12^.

Propagation-based phase contrast is one of the simplest approaches for generating phase-contrast images because it does not require additional optical elements. In this technique, a spatially coherent X-ray beam is required. The beam propagates through the sample where induced phase shifts result in refraction of the X-ray beam. The detector is positioned at a distance from the sample, allowing the phase variations to be converted into detectable intensity variations. In the hard X-ray range (photon energy above 10 keV), the contribution from phase contrast can be three orders of magnitude higher than that from absorption contrast^11^.

In magnetic resonance imaging (MRI), improving cancer assessment is a major focus, particularly through the development of diffusion-weighted MRI (DWI) methods capable of detecting the microstructural tissue changes associated with cancer detection and grading^13^. DWI is a natural choice for cancer assessment as the microstructure changes that are used to define and characterise solid tissue cancers through histopathology are thought to strongly determine the water mobility characteristics that affect DWI signal intensity. DWI based “microstructure imaging” techniques aim to infer sub-voxel tissue structure by analysing how the tissue microstructure interacts with the measurement process to produce the detected MRI signal. See^14,15^ for prostate specific overviews.

In the case of prostate cancer assessment, there is particular interest in developing imaging biomarkers that correlate with Gleason grading, as these histological features are critical for prognosis and treatment decisions. Gleason patterns are used to describe the architectural appearance of prostate cancer tissue as seen in light microscopy of stained sections. They are part of the Gleason grading system^16^, which is widely used to assess the aggressiveness of prostate cancer and consequent therapy and management decisions. The system classifies tumour gland patterns on a scale from 1 to 5. Gleason patterns 2-3 represent well-formed cancer glands that still resemble normal prostate glands but are abnormal in their arrangement. Gleason pattern 4 shows loss of normal glandular structure and more complex tumour architecture. pattern 5 describes complete loss of gland structure with heightened risk of metastatic disease. In current clinical practice, patterns 1 and 2 are regarded as benign. The Gleason grade is based on reporting the most common, and second most common Gleason pattern observed in a tissue sample. “Clinically significant” cancer (requiring monitoring or therapy intervention) is generally defined as a predominance of patterns 4 or 5. Thus microstructure imaging methods are often assessed according to their ability to discriminate benign tissue and pattern 3 from patterns 4 and 5.

Although promising, the development and optimisation of MRI-based microstructure prediction methods remain limited by validation approaches that rely primarily on two-dimensional histology. A recent study addressed this problem by using MRI microscopy performed at 20 *µ*m resolution to estimate relative volumes of epithelium, lumen, and stroma components contributing to the DWI signal measured at 160 *µ*m resolution^17^. Nevertheless, the spatial resolution of this MRI microscopy remains well above the typical water diffusion length scale probed by the DWI method^18^, and thus provides only a meso-scale average of larger gland architecture features without true microstructure detail.

Three-dimensional *µ*CT may offer particular value for DWI validation, as it has the potential to provide relevant microstructural information across the full volume corresponding to each MRI voxel, at a spatial scale that represents structure features likely defining the water displacements detected by DWI methods. A study of mouse brain^19^ provides an example wherein *µ*CT derived neural fibre orientation was demonstrated to be strongly correlated with MRI derived fibre tract orientation.

A recent *µ*CT study of formalin fixed paraffin embedded (FFPE) prostate needle biopsy samples demonstrated the potential for *µ*CT to provide 3D structural information similar to standard histology and light microscopy, with the possible novel advantage of permitting multi-level and alternate plane views that can discriminate subtle differences between some benign and malignant gland features^12^. This example addresses the fundamental undersampling problem of thin section approaches to tissue structure characterisation.

In this paper we present results from a *µ*CT study of larger, 3 mm diameter, samples of human prostate tissue obtained from radical prostatectomy specimens, and imaged in an 80% ethanol matrix. Results are compared with post *µ*CT histology of the samples. In addition illustrate the potential for *µ*CT-based validation of DWI methods based on a fibre orientation analysis of the *µ*CT data.

## Results

Fig. 1 gives an overview of typical *µ*CT results for a sample of normal (non-cancerous) glandular prostate tissue. An aligned H&E stained thin section of the same sample demonstrates the capacity of *µ*CT to provide 3D structure information not available from the histological section, albeit at a lower spatial resolution. *µ*CT provides excellent soft tissue contrast, clearly delineating normal gland architecture with distinct contrast between the acellular lumen space and the cellular epithelium and stroma.

**Figure 1.**
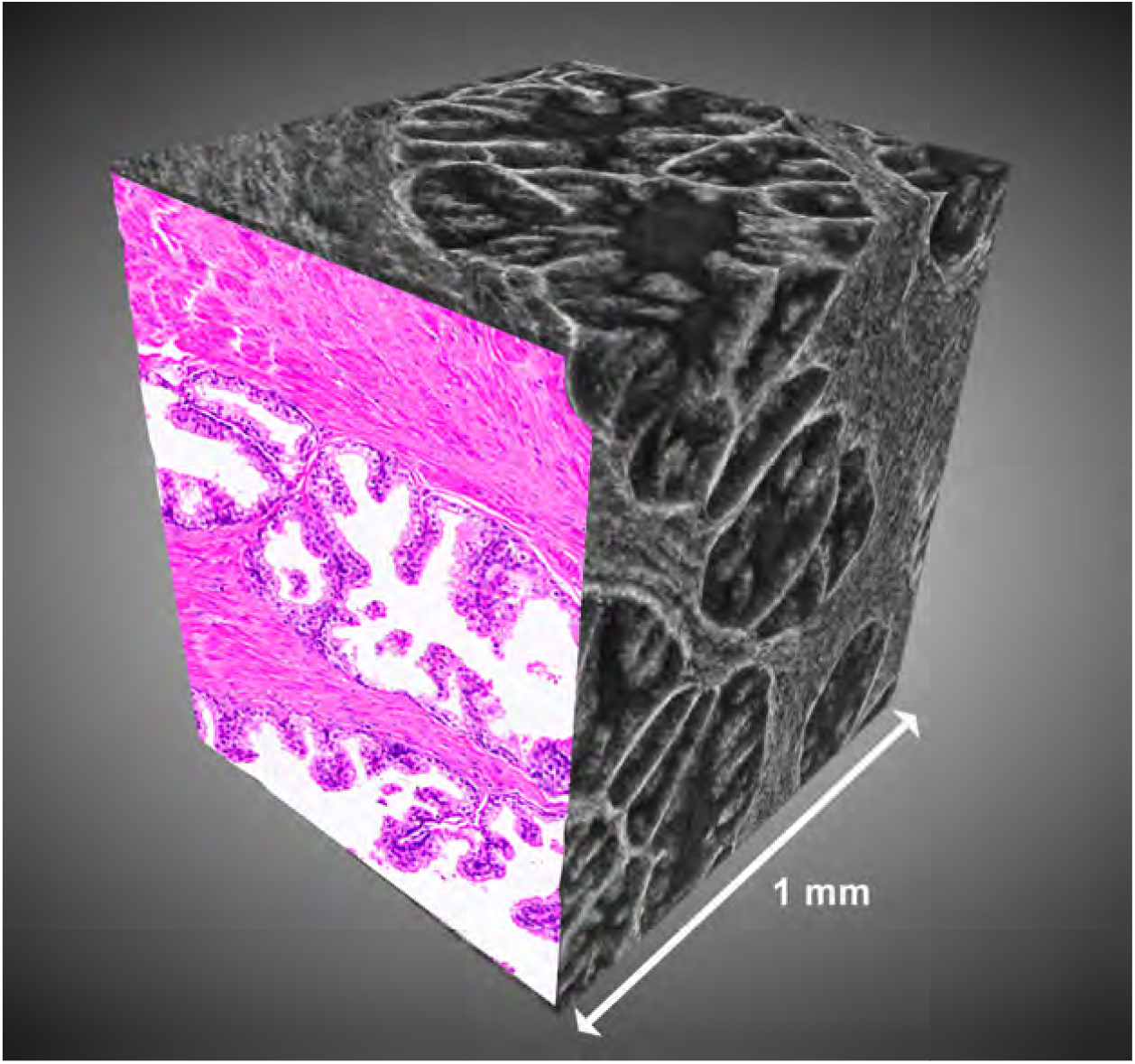
Typical *µ*CT results for a sample of normal glandular prostate tissue. The 3D *µ*CT volume with aligned histology section demonstrates the extensive extra microstructure information available from *µ*CT. Reconstruction of the equivalent volume from histology would require microtomy and staining of ca. 200 serial sections. Labelled gland components are shown in Fig. 2. Sample from Patient 1.

Fig. 2 shows detail of normal gland acini and the corresponding H&E stained section. The ca. 3 *µ*m spatial resolution of the *µ*CT enables resolution of some subcellular components, although not sufficiently to allow unequivocal identification. Based on^12^ the larger dark regions in the epithelium are likely cell nuclei (see Discussion).

**Figure 2.**
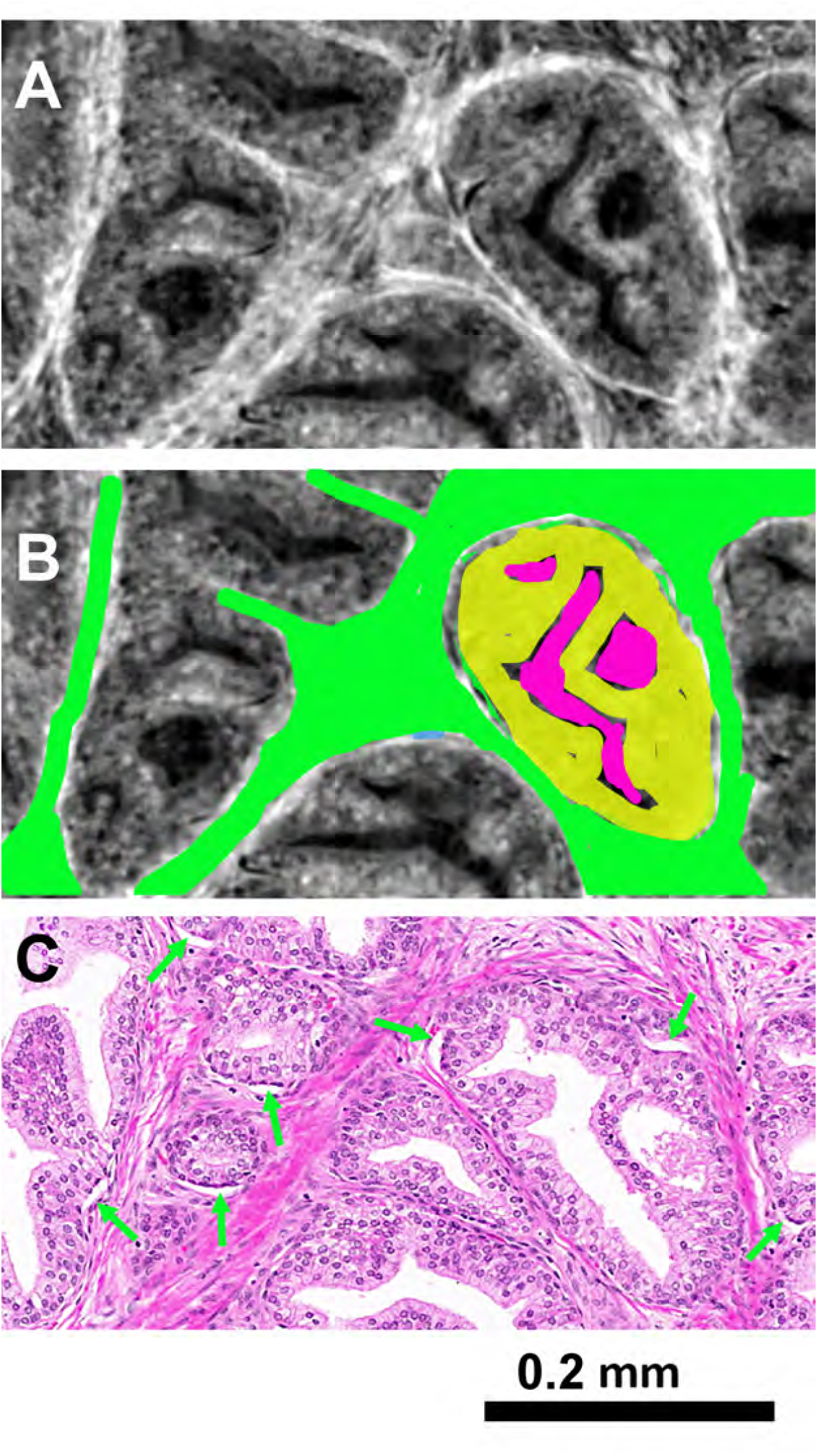
Glandular architecture and microstructure detail. A) Normal glanudular tissue *µ*CT section. B) *µ*CT with colour overlay defining typical gland components. A gland acinus is comprised of secretory columnar epithelial cells (yellow) surrounding a lumen space (pink). Note that only one gland acinus has been coloured. The gland acini are embedded in a fibromuscular stromal matrix (green). C) Corresponding approximately aligned H&E stained histology section. The green arrows show examples of epithelial layer detachment from the basement membrane. This detachment (attibutable to paraffin embedding) is absent in the corresponding *µ*CT section (see Discussion). Sample from Patient 6.

In normal prostate glands the columnar epithelium has a thickness of around 40-50 *µ*m, with typical nuclei around 6-9 *µ*m diameter. Based on size and location, it is probable that the low X-ray density (dark) round regions in the epithelium represent nuclei.

In the histology section of Fig. 2C there are instances of the epithelial layer detaching from the basement membrane. This common histology processing artifact, attributed to the paraffin enbedding step, is rarely seen in the mCT images.

Fig. 3 shows a sample of benign glandular tissue with bands of fibromuscular stroma. This sample includes examples of corpora amylacia – solid glycopolymer particles commonly seen in prostate tissue but not having diagnostic value. Some of the corpora amylacia show layering (‘onion rings’) on both *µ*CT and the H&E section. Movement artifact (a double ‘crescent’) can be seen on some particles, suggesting they may be mobile in the liquid and largely acellular gland acinus space.

**Figure 3.**
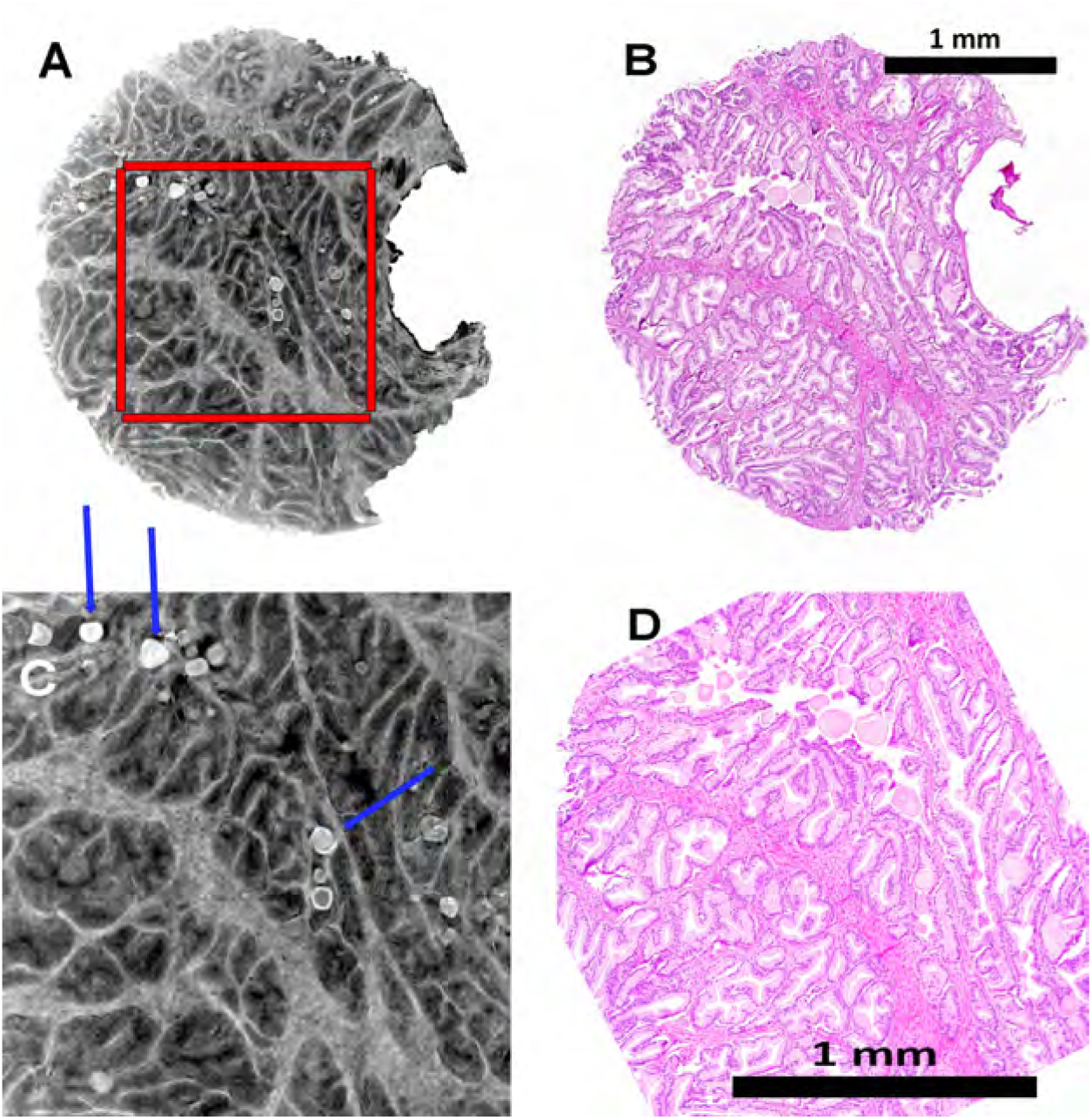
Benign glandular tissue. A, B) Full section. The red square marks the location of detail shown in C and D. Blue arrows show examples of corpora amylacia – solid particles primarily composed of glycopolymers, often often having a layered ‘onion rings’ appearance. There is evidence of movement artifact in some corpora amylacia in the *µ*CT section (rightmost blue arrow). The H&E-stained histology section shows the closest match to the *µ*CT section. Sample from Patient 3.

Fig. 4 shows a further example of benign glandular tissue, inthis case with large bands of distinctly fibrous stroma. The orientation of the apparent fibres in the *µ*CT is consistent with the orientation of cells and extracellular fibre in the H&E stained section. This sample, and those shown in Figs 1 and 2, show a distinct narrow bright band around each gland acinus in the *µ*CT images – most likely corresponding to the protein-dense basement membrane. The basement membrane is not normally evident on H&E staining, and requires specific additional histochemical stains for visualisation on light microscopy. This layer is much less obvious in Fig. 3, possibly evidence of a patient-specific structural variation.

**Figure 4.**
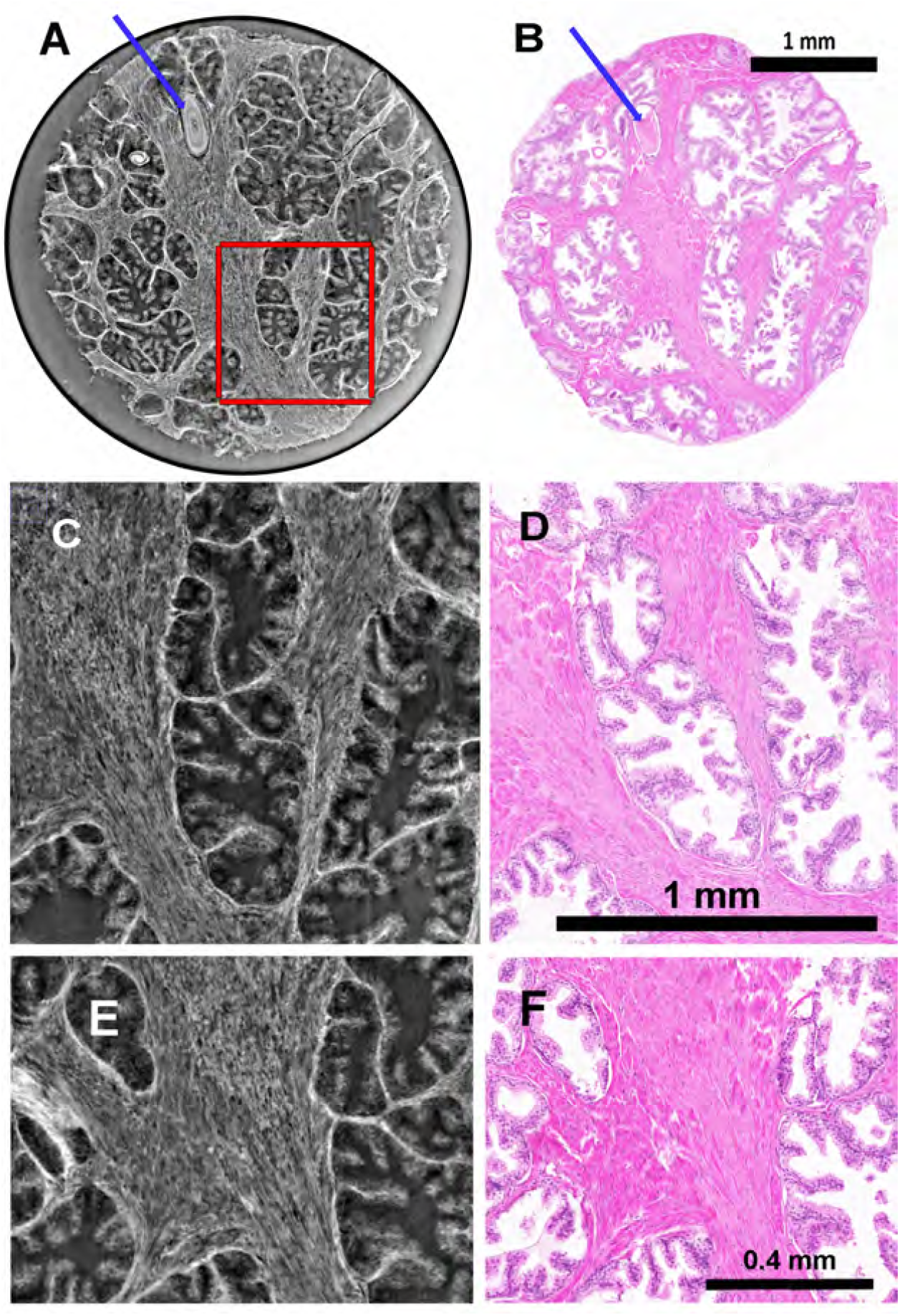
Benign glandular tissue with distinctly fibrous stroma. A, B) Full section. The red square marks the location of detail shown in C and D. Blue arrows show a large corpus amylaceum with typical ‘onion rings’ layered structure evident in the *µ*CT image. E, F) Detail illustrating close correspondence between stromal fibre orientations in *µ*CT and histology section. Sample from Patient 4.

Fig. 5 shows a further variant of mostly normal glandular tissue, in this case with distinct papillary epithelium. The H&E section shows a small focus of adenocarcinoma (too small for assignment of a Gleason pattern) that is notably not evident as a distinct structural variation on the corresponding *µ*CT.

**Figure 5.**
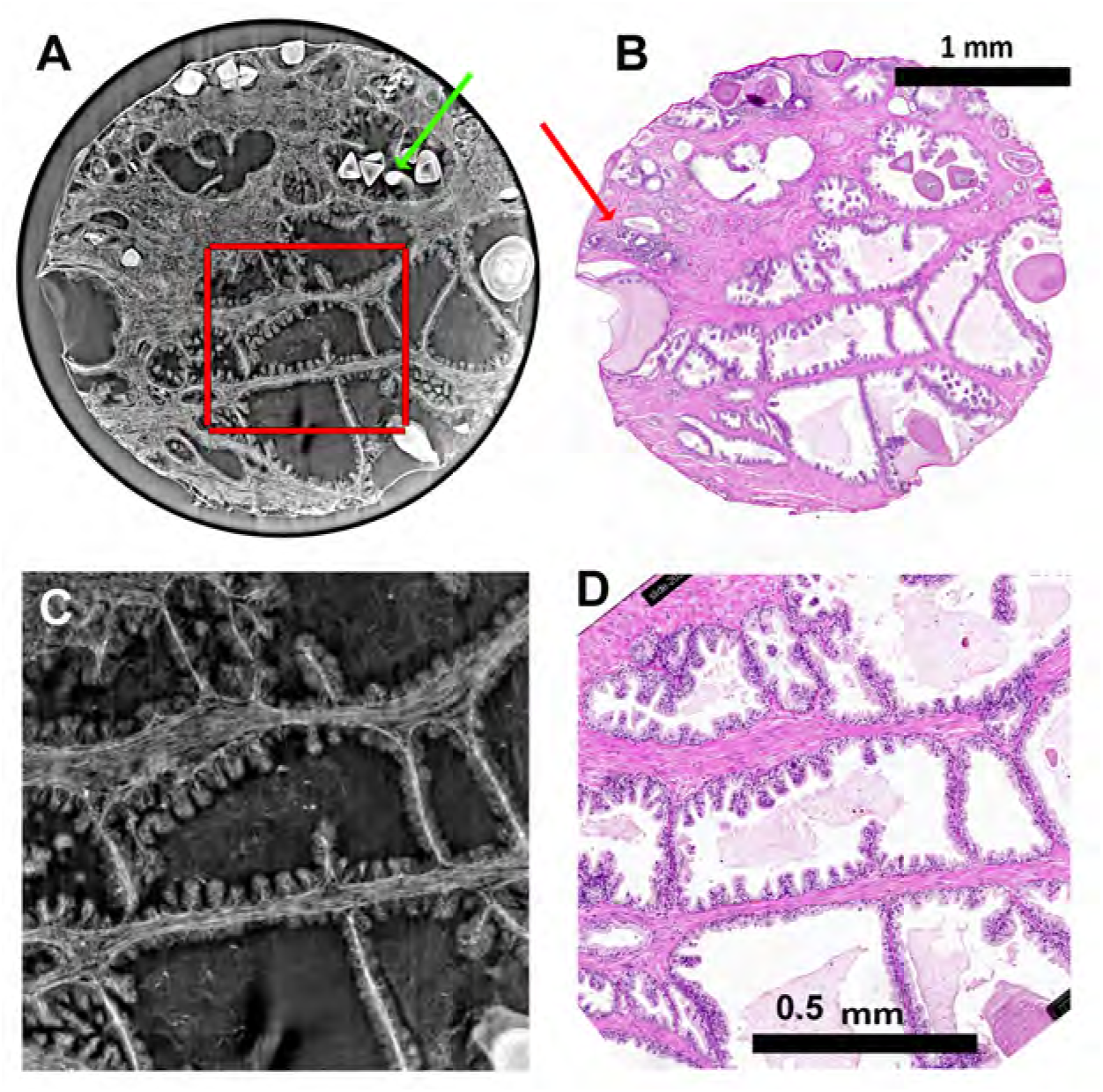
Benign glandular tissue with papillary epithelium. A, B) Full section. The red square marks the location of detail shown in C and D. The H&E section shows a 0.4 ×0.2 mm microfocus of adenocarcinoma (red arrow), not evident in the corresponding *µ*CT section. The green arrow shows evidence of movement artifact in some corpora amylacia. Sample from Patient 2.

Fig. 6 shows a sample containing regions of Gleason pattern 3 and pattern 4 cancer. The pattern 4 cancer is lobular (large clumps of tightly packed gland acini) and has a cribriform structure where proliferating epithelial cells form bridges across the luminal space. The adjacent pattern 3 cancer has highly disordered glands in a non-lobular arrangement. In both cases the different tissue architectures are clearly evident in the *µ*CT and correspond closely to the structure seen in the H&E section.

**Figure 6.**
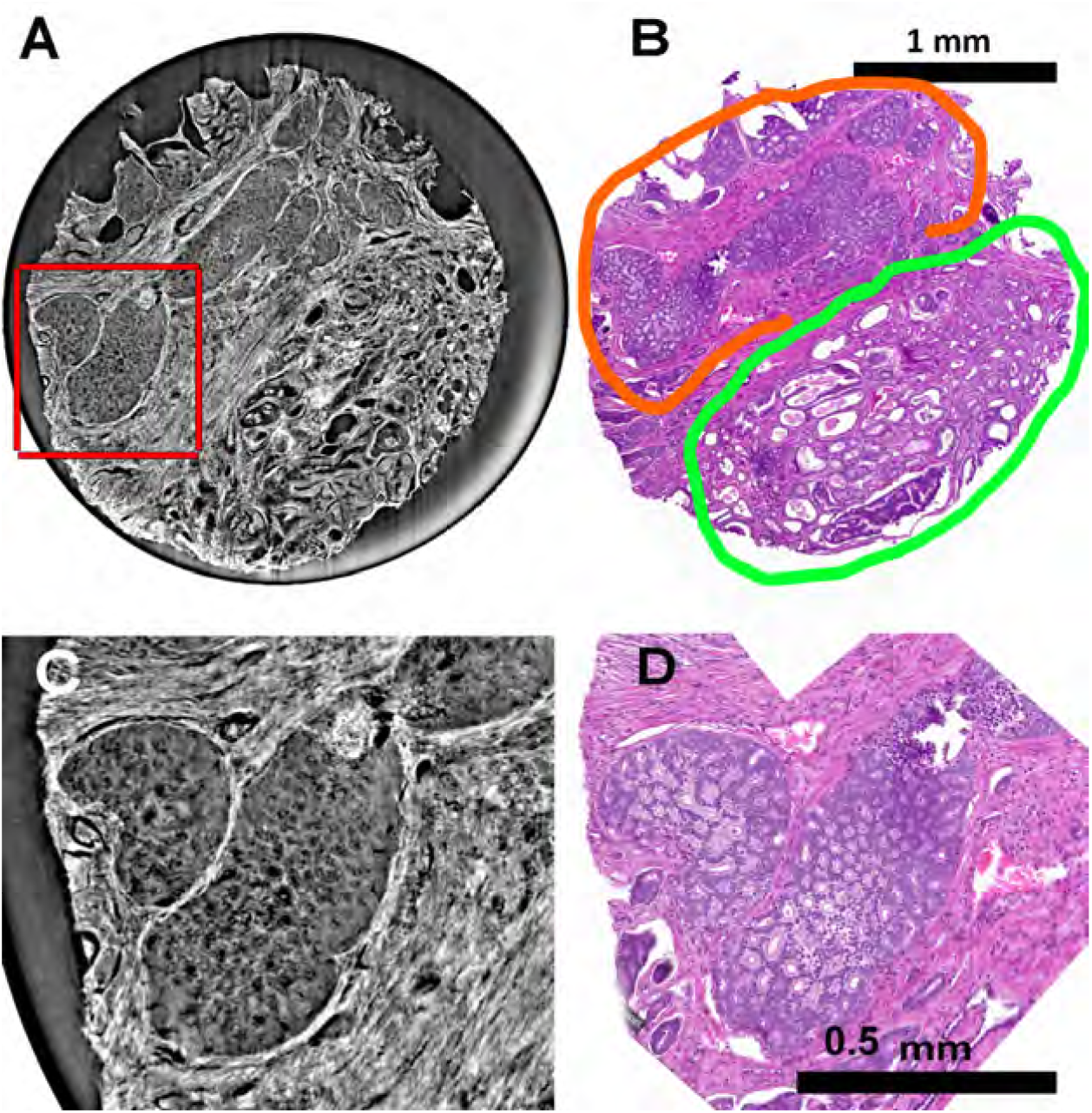
Cancer sample. Gleason patterns 3 and 4. A, B) Full section. The red square marks the location of detail shown in C and D. Orange outline shows adenocarcinoma with cribriform Gleason pattern 4. Green outline shows Gleason pattern 3 with disordered non-lobular arrangement. Sample from Patient 7.

Fig. 7 shows another example of cancer with Gleason patterns 3 and 4. Although distinctly different in architecture from the sample shown in Fig. 6 *µ*CT again shows the major structural differences evident in the H&E section.

**Figure 7.**
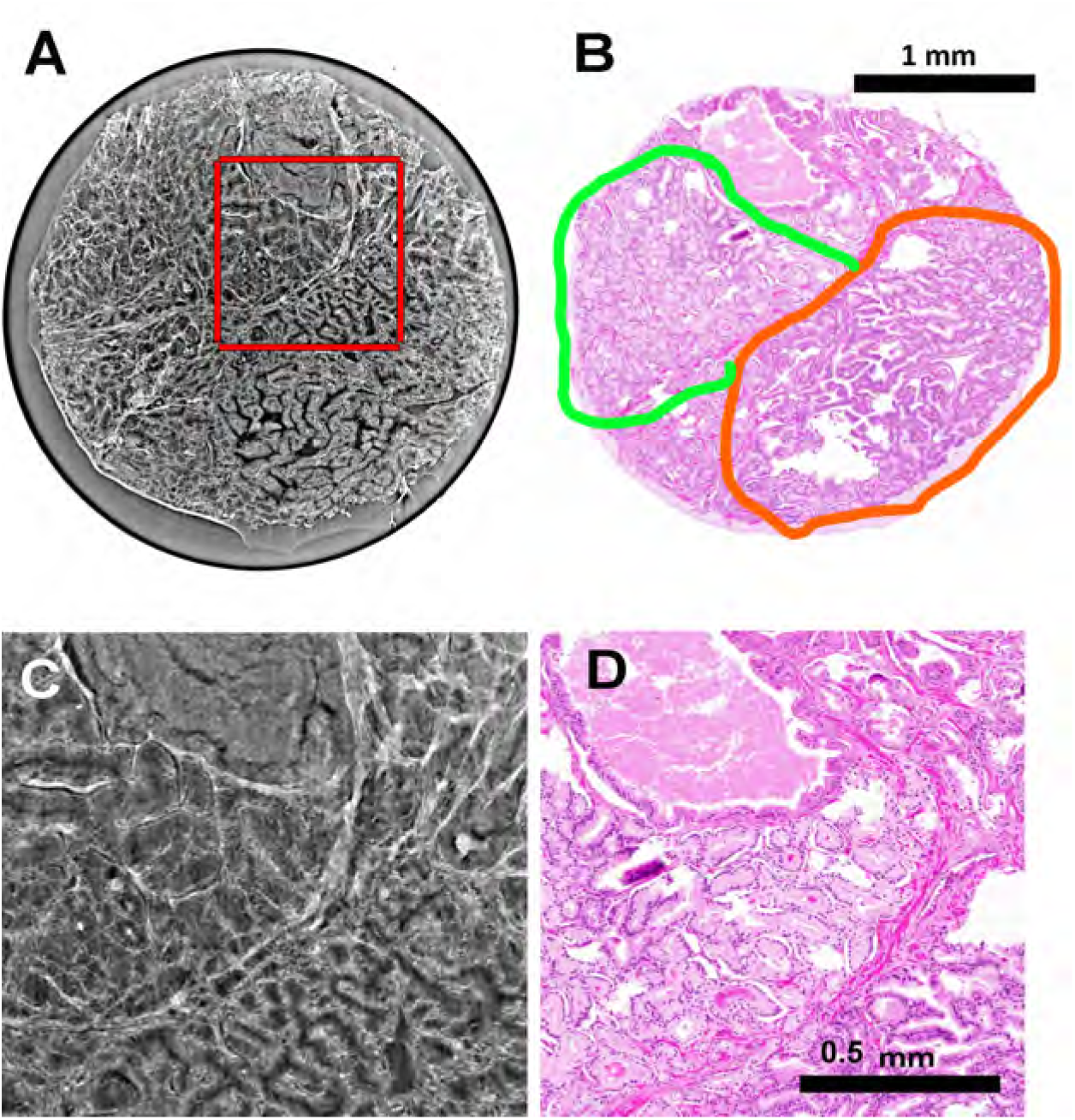
Cancer sample. Gleason patterns 3 and 4. A, B) Full section. The red square marks the location of detail shown in C and D. Images show an interface between two tumour morphologies, Gleason patterns 3 (green outline) and 4 (orange outline), with high density of abnormal glands, non-lobular in patterning, and small lumina. Sample from Patient 5.

Fig. 8 shows results from DWI microscopy and subsequent *µ*CT of a sample of normal glandular tissue (sample as for Fig. 4). *µ*CT fibre orientation analysis gave results consistent with the visually apparent stromal fibre direction, and in broad agreement with the DWI-derived primary diffusion direction and fibre tracking. It should be noted that in this preliminary example the *µ*CT based fibre orientation analysis has not been fine-tuned to match the spatial scale of water displacements and spatial averaging that define the DWI signal intensity, so a close match between images B and D is not expected.

**Figure 8.**
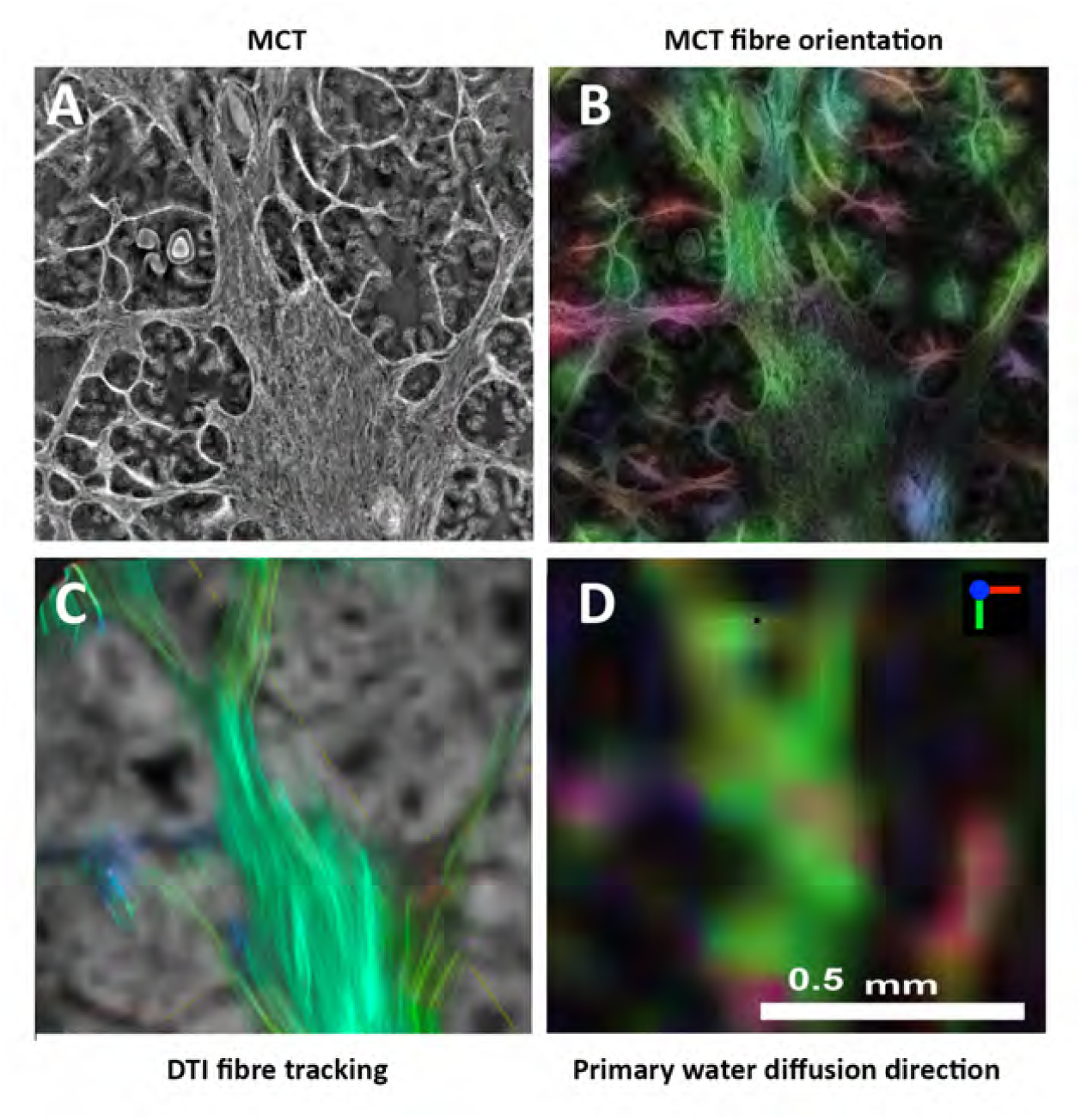
Comparison of *µ*CT and magnetic resonance diffusion weighted imaging (DWI). A) *µ*CT of normal glandular tissue showing glands, corpora amylacia, and a large central area of fibromuscular stroma with the majority of fibres running top to bottom of the image. This image shows one slice of the stack of 28 *µ*CT slices that lie within the thickness of the corresponding MRI slice. B) *µ*CT-derived fibre orientation. Colour-coded direction as shown in (D) inset. The colour intensity is scaled by a confidence metric to suppress non-fibrous regions^20^ so that glands, lumen space, and corpora amylacia show dark or black. The fibre orientation is based on analysis of a stack of slices centred on that shown in image A. C) DWI background (voxel size 40 × 40 × 40 *µ*m) with overlayed fibre tracking based on DWI-derived preferential water diffusion direction. D) DWI-derived preferential diffusion direction at slice corresponding to images A–C. Note the qualitative concordance between DWI- and *µ*CT-derived fibre orientation. Sample from Patient 4.

## Discussion

This paper presents a survey of synchrotron phase contrast *µ*CT of a small range of prostate tissue samples with corresponding histology sections of the same tissue. We also present a demonstration of the potential of *µ*CT to provide a 3D microstructure information for testing and validation of diffusion-weighted MRI.

### Sample preparation for *µ*CT

Our gelatin embedding technique (based on^21^) enabled a stack of 2–3 tissue cores to be imaged in a water matrix for MRI, and then transferred to ethanol for *µ*CT without disturbance of tissue orientation. With the tissue samples fixed inside a plastic tube that was a close fit inside a standard 5-mm nuclear magnetic resonance sample tube (Fig. 9) the Z-axis of the MRI scan was collinear with the axis of rotation in *µ*CT when the same plastic tube was mounted in the 3-jaw chuck of the *µ*CT stage. This arrangement simplifies alignment of MRI and *µ*CT data for downstream correlation analysis. Direct transfer of the embedded samples from water to 80% ethanol resulted in conspicuous shrinkage of the gelatin^22^. This was avoided by stepwise transfer via 40 and 60% concentrations, with ca. 24 hr in each.

**Figure 9.**
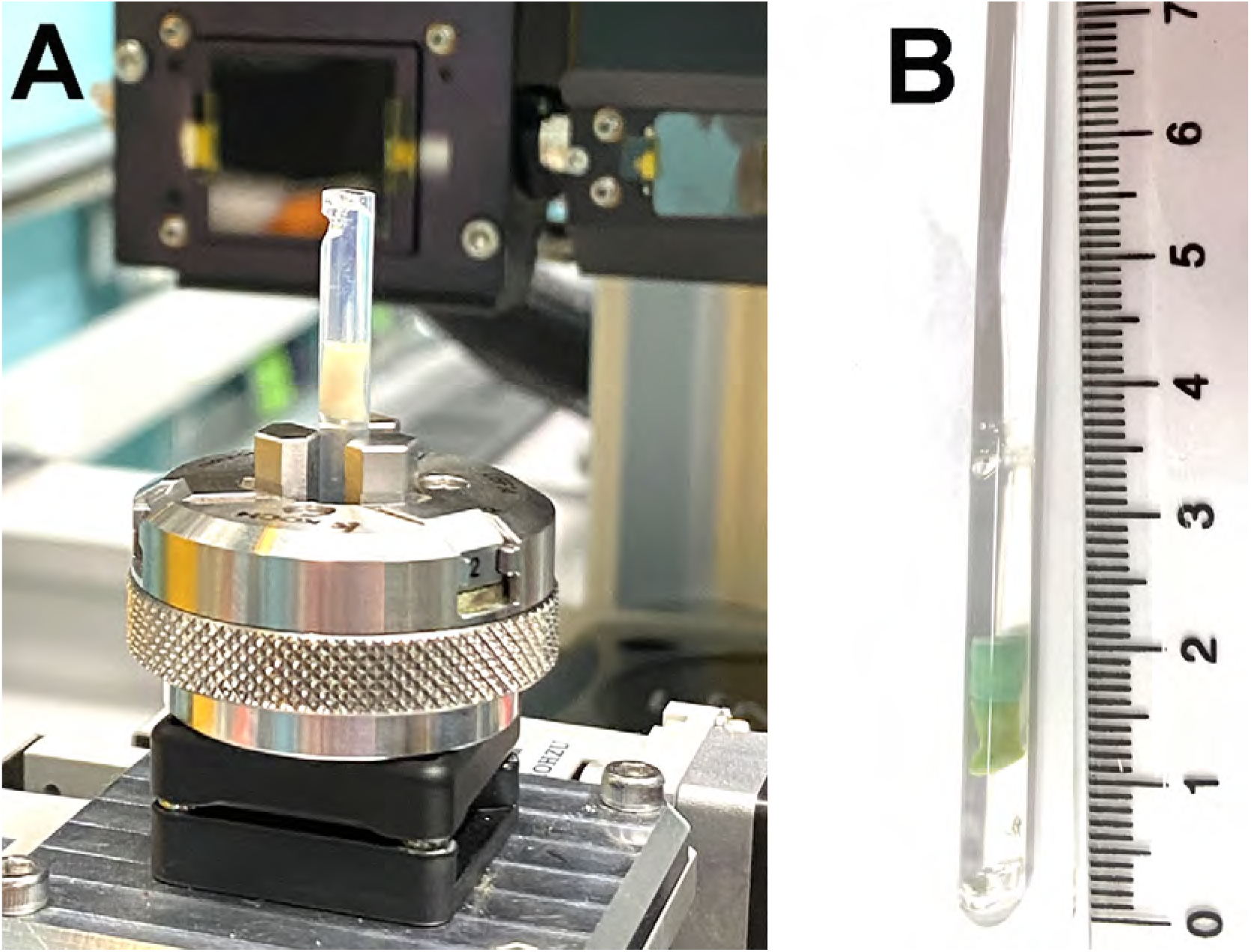
Sample mounting for *µ*CT. A) Gelatin-embedded tissue core in PTFE tube mounted on *µ*CT stage. B) Embedded samples in PTFE tube can be inserted into a 5-mm NMR (nuclear magnetic resonance) tube to acquire magnetic resonance imaging data in precise alignment with *µ*CT (See Fig.8).

Sample movement and gas bubble formation are reported as common problems in soft tissue *µ*CT^21,23^, and contrast enhancement with an ethanol matrix may lead to tissue shrinkage^24^. We saw no clear evidence of tissue shrinkage due to the 80% ethanol *µ*CT imaging matrix. Notwithstanding the spatial resolution differences, tissue sample size relative to the enclosing plastic tube was the same in MRI and *µ*CT views at matched locations. Histological processing for light microscopy introduced multiple additional tissue distortions and is not a useful gauge for shrinkage post fixation shrinkage due to ethanol. We also found no evidence of movement of the sample, although there were apparent movement artifacts around some corpora amylacia. Although we used a continuous *µ*CT stage rotation, expected to be free from mechanical stepping vibrations, many of the smaller corpora amylacia would be free to move in the surrounding liquid-filled gland lumen space – possibly due to thermal heating from the X-ray beam. Motion artefacts can be minimised by reducing the exposure time because the sample has less time to move while the detector is collecting the image, however, this will decrease the signal-to-noise ratio in the reconstructed images. Given the lack of structural importance of corpora amylacia, any movement artifacts are probably of no significance to most *µ*CT applications.

We degassed all liquid solutions under vacuum in order to minimise the chance of gas bubble formation during sample handling or imaging, as per^21^. No bubble formation was observed, even in samples scanned multiple times in succession. It is possible that in our setup any heating effects were dissipated by the mass of the plastic tubes in which the gelatin-embedded samples were imaged.

### Tissue microstructure information available from *µ*CT and comparison with conventional histology

Our *µ*CT images of tissue in an 80% ethanol matrix showed distinctly higher contrast than those previously presented for dog prostate tissue in a water matrix^11^, and much higher contrast and spatial resolution that a previous report of phase contrast projection imaging of a 3-mm thick slice of human prostate tissue in a water matrix^3^. This contrast enhancement is likely due to the lower X-ray density of ethanol relative to water, in turn due to the effective replacement of oxygen with carbon.

Our tissue contrast was also very different from that reported for unstained formalin fixed paraffin embedded (FFPE) human prostate biopsies^12^. In contrast to 80% ethanol versus water, the X-ray attenuation difference between 80% ethanol and paraffin should be relatively small as each have similar carbon/hydrogen density. Rather, the contrast discrepancy may reflect different photon energy (14 keV in^12^ versus 22 keV in our study), and propagation distance (50 mm in^12^ versus 155 mm in our study). Either way, these differences highlight the potential dependence of contrast and spatial resolution on the beamline hardware of the specific synchrotron.

The spatial resolution (ca. 3 *µ*m) and contrast of our *µ*CT method provides prostate tissue microstructure information on an approximately cellular scale, but does not provide the subcellular detail available from conventional histology with H&E staining and light microscopy. The study by Sugarman et al^12^, with a spatial resolution of ca. 1 *µ*m and precise alignment of *µ*CT and histology sections, was able to unequivocally identify epithelial cell nuclei, but not smaller diagnostically important features such as nucleoli. In our study we saw dark, roughly circular, regions in the epithelial layer of the *µ*CT, however, as we did not have precisely aligned histology sections we are unable to confidently identify these as nuclei.

Several of the presented H&E stained sections show detachment of the epithelial layer from the basement membrane (Labelled examples in Fig. 3C). This detachment is not seen in the corresponding *µ*CT images. This detachment is a know artifact attributed to the paraffin embedding step^25,26^, and illustrates a potential advantage of *µ*CT over histology processing for some investigations.

Aside from the non-destructive provision of 3D structural information, *µ*CT has other potential advantages over histology, depending on the information required. Although not resolved, questions over the reliability of histology-based measurements of epithelium, stroma, and lumen volumes may affect validation of MRI “microstructure imaging” methods that specifically model and predict changes in the relative volumes of these components, eg.^27,28^.

The mostly thin hyperintense (white) layer around gland acini seen in many of our normal tissue images is most likely the collagen fibre dense basement membrane^29^. There is no corresponding contrast in the H&E sections, consistent with the requirement for extra chemical or immuno stains for visualisation of the basement membrane on light microscopy. Similar collagen hyperintensity is seen in *µ*CT of breast tissue^5^, although notably not in the FFPE prostate tissue *µ*CT study^12^.

### Microstructural validation of diffusion MRI methods

The 3D microstructure information available from phase contrast *µ*CT is potentially invaluable for validation and development of MRI methods, particularly for diffusion-weighted imaging (DWI) where the biophysical contrast mechanism depends directly on the way tissue microstructure modifies the mobility of water molecules^30^. Correlation with post-surgery histology and light microscopy is the current gold standard for validation of clinical MRI techniques that aim to characterise tissue structure, but is compromised by multiple technical problems^31^, not least being the inability of 2D histology to fully represent the 3D tissue structure that determines DWI contrast and signal intensity. Phase contrast CT (at both micro and meso scales), if reliably aligned with MRI data, opens the possibility of an unprecedented voxelwise correlation of MRI with underlying structural information. In our example (Fig. 8), each DWI voxel (40 *µ*m isotropic) corresponds to ca. 2400 *µ*CT voxels (effective spatial resolution ca. 3 *µ*m isotropic), enabling a wealth of structural detail to inform the observed signal differences between DWI voxels. A recent demonstration of whole prostate synchrotron phase contrast CT suggests the possibility of a direct correlation with in vivo prostate MRI^32^, although it should be noted that reregistration of imaging data acquired in vivo with imaging data from a resected organ remains a major technical challenge^31^.

Our sample mounting method has enabled a direct comparison of closely aligned DWI and *µ*CT data, and demonstrates the ability of *µ*CT to provide fibre orientation information that supports conclusions drawn from diffusion tensor analysis of DWI data. This is the first (albeit qualitative and preliminary) 3D structure comparison of the diffusion modifying effects of fibromuscular stroma on water diffusion measured by MRI – a major advance on the limitations of a previous 2D histology correlation^33^. A quantitative DWI-*µ*CT correlation analysis is beyond the scope of this preliminary investigation and will require, in addition to an extensive sample range, optimisation of the Gaussian smoothing steps of the *µ*CT fibre analysis to match the MRI diffusion time and voxel size^20^, and a more sophisticated DWI method than that employed in the current study.

The potential of *µ*CT-based validation of DWI is not limited to fibre orientation analysis. *µ*CT-derived porosity and tortuosity metrics may provide structural validation of apparent diffusion coefficient (ADC) measurements from DWI, and extend our understanding of the diffusion time dependence of ADC measurements in soft tissues^18^. A recent study of monkey brain tissue used *µ*CT for 3D validation of DWI-based measurements of axonal morphology^34^.

### Study limitations

This study presents a preliminary survey of the capabilities of *µ*CT for characterisation of human prostate tissue. We performed a qualitative optimisation of the scan parameters based on the facilities available at the Australian Synchrotron. We found distinct differences from previous phase contrast CT studies of prostate tissue, highlighting the likely site-specific variations to be expected from this technique. Our results are specific to the sample preparation method and beamline setup we describe. Both spatial resolution and image contrast would be expected to be different for alternative sample preparations and imaging methods, which would need to be optimised according to the specific research questions addressed.

For many studies, particularly those aiming for precise histology correlation, scanning of paraffin embedded samples may be more appropriate, and would result in different contrast characteristics from those we observed.

Phase contrast *µ*CT, of the spatial resolution and quality described here, requires an intense coherent monochromatic light source, currently only available from synchrotrons. This limits the availability of the technique, although the development of compact ‘laboratory’ light sources is expected to broaden future access to the technique.

## Methods

### Tissue samples

All tissue samples were collected with written and informed patient consent under approval by the Sydney Local Health District Research Governance Office. (Project ETH00217). Samples and data were handled in accordance with the Australian National Health and Medical Research Council guidelines.

Samples (Table 1 were taken from seven formalin-fixed radical prostatectomy specimens during routine processing for histopathology reporting. All patients For each prostate, a 3-mm Stiefel core punch was used to cut 2-4 tissue cylinders from the 4-5 mm thick macro sections. Sampling sites were selected by an expert histopathologist with the aim to provide a diversity of tissue types, both malignant and benign, at locations which would not affect the clinical reporting results. Note that at the time of sampling unequivocal identification of cancerous and normal tissue is generally not possible. The actual pathological status of each sample was assessed by standard histological processing and light microscopy examination at the conclusion of the study. Samples were stored in neutral buffered formalin at room temperature. Storage times varied from 1-4 months depending on available synchrotron beamtime.

**Table 1.** Samples and patients.

| Patient | Sample pathology | Figure | Pre-mCT MRI | mCT projections |
| --- | --- | --- | --- | --- |
| 1 | normal | 1 | no | 1800 |
| 2 | normal | 5 | yes | 3600 |
| 3 | normal | 3 | yes | 3600 |
| 4 | normal | 4, 8 | yes | 3600 |
| 5 | Gleason patterns 3 & 4 | 7 | yes | 3600 |
| 6 | normal | 2 | no | 3600 |
| 7 | Gleason patterns 3 & 4 | 6 | yes | 3600 |

### Sample preparation

Fixed tissue cores (3 mm diameter by 4-5 mm long) were degassed at room temperature in formalin under vacuum for one hour and then inserted into a 4 mm OD 30 mm long PTFE (polytetrafluoroethylene) tube for gelatin embedding. A 5% gelatin solution was made up in hot degassed milli-Q water (McKenzies crystal gelatin, pork origin, Ward McKenzie Pty Ltd, Victoria, Australia). After cooling to ca. 40°C, the gelatin was injected with a syringe and needle into the space around the tissue cylinder, ensuring ejection of any air bubbles and filling of the PTFE tube. The PTFE tube was then laid horizontally, immersed in liquid gelatin, and allowed to set overnight at room temperature. After setting of the gelatin, the PTFE tubes were extracted and transferred to 80% ethanol in milli-Q water via overnight steps in 40%, and 60% ethanol. Direct transfer to high ethanol concentrations resulted in severe gelatin shrinkage. Gelatin-embedded samples were stored in 80% ethanol for 1-2 months at room temperature prior to *µ*CT. We note that possible osmotic effects were inherent in the applied sample treatment protocol. Post *µ*CT histology (as assessed by an expert prostate histopathologist) showed no evidence of overfixation or tissue degradation due to osmotic shock or bacterial contamination during sample storage.

### Propagation-based phase contrast *µ*CT

Propagation-based phase-contrast *µ*CT was performed in two beamtime sessions on the micro computed tomography beamline at The Australian Synchrotron, Melbourne, Victoria. The *µ*CT beamline is a 1.3 T bending-magnet beamline on a third-generation 3 GeV (200 mA) synchrotron light source. Monochromatic X-ray beams with approximately 3% bandwidth are produced using a double multilayer monochromator.

Gelatin-embedded samples in PTFE tubes were mounted vertically in a 3-jaw chuck on the *µ*CT stage with the bottom of the tissue cylinder 1-2 mm above the top of the chuck jaws (Fig. 9). To ensure sample rotation around the axis of the sample tube the sample tube ends were not sealed or capped. There was no significant loss of fluid (80% ethanol) from the gelatin embedding medium during the ca. 10 min *µ*CT scan. There was no evidence of gas bubble formation during the *µ*CT scan^23^. Post-*µ*CT, samples (gelatin-embedded in PTFE tubes) were stored in 80% ethanol for 1-2 weeks prior to histological processing.

*µ*CT imaging and reconstruction parameters (photon energy, propagation distance, *γ*) were optimised to produce a subjectively assessed maximum image quality in terms of visibility of assumed glandular, stromal fibre, and subcellular structure details. Tested ranges included photon energies 15-28 keV, propagation distances 15.5-20 cm, and *γ* range 100-1000 based on previous experiments on this beamline^35^. 15.5 cm is the shortest propagation distance possible on this beamline, with image blurring increasing with longer distances. Photon energies less than ca. 17 keV resulted in poor X-ray transmission, and energies greater than 25 keV led to reduced tissue contrast. The selected *γ* produced a balance between edge sharpness and contrast detail that enabled the most subjectively clear distinction of microstructure details.

After optimisation the following imaging protocol was adopted: 22 keV monochromatic beam with 3% bandwidth; 1800 or 3600 projections with 180 degrees sample rotation; sample to detector propagation distance 15.5 cm. A white beam detector (PCO.edge 5.5 cMOS camera, 35 *µ*m LuAG:Ce scintillator) was used with 4.5X objective lens giving a field of view (FOV) of 3.7mm horizontal by 3.1 mm vertical and effective pixel size of 1.44 *µ*m with image shape 2560 by 2160 pixels. Exposure time was 70 ms per projection for a scan time of ca. 10 minutes per sample. In the fly scan mode used the sample rotates continuously while projection data are captured by the detector. In this mode, the scan time is determined by the rotation speed of the stage rather than by the number of projections. Before the scan 100 dark (beam off) and 100 flat field (beam on, no sample) images were acquired for correction of projection data. Spatial resolution was ca. 2.7 *µ*m in the transaxial plane and ca. 2.5 *µ*m in the sample axis direction, based on FWHM of the point spread function^35^. The change from 1800 projections (beamtime session 1) to 3600 projections (beamtime session 2) was based on an observation of slightly better image quality with the larger number of projections. Only the sample from Patient 1 (Fig. 1) was imaged with 1800 projections.

Data were reconstructed using the *µ*CT processing pipeline^35^. Projection data were corrected using flat-field and dark-field images to account for imaging system non-uniformities. Transport of Intensity Equation (TIE)-based phase retrieval^36^ was applied to each projection using *γ* = 244. Three dimensional reconstruction was then performed on the phase-retrieved projection images^37^. Total pipeline processing time was ca. 5 minutes. FIJI^38^ and ORS Dragonfly (Comet Technologies Canada Inc. Dragonfly 3D World Version 2025.1. https://dragonfly.comet.tech/) software were used for 2D and 3D visualisations respectively.

Fibre orientation analysis of *µ*CT data was performed by 3D structure tensor analysis^20^ using in-house Matlab code. Briefly, the method takes a stack of *µ*CT slices (corresponding to the matched DWI slice) and calculates a 3D image gradient at each *µ*CT voxel. The direction of the minimum gradient is taken as the fibre direction. A line-weighted fractional anisotropy was used as a weighting to minimise the effects of hard edges (eg. of corpora amylacia). An initial 3D Gaussian smoothing of the *µ*CT data corresponds to the diffusion time parameter in a DWI measurement, and a final smoothing emulates intra-voxel signal averaging.

### Histological processing and analysis

Prior to standard histology processing (Charles Perkins Centre, The University of Sydney), the tissue cylinders were gently ejected from the PTFE tubes and excess gelatin trimmed from the ends with a scalpel blade. Samples were then processed and paraffin-embedded with the axis of the tissue cylinder vertical in the block so that subsequent microtomy would produce sections in a similar orientation to the transverse *µ*CT image plane (orthogonal to the tissue cylinder axis). 5-10 5-*µ*m sections were acquired at ca. 0.3 mm intervals through the depth of the tissue block to ensure multiple histology sections would be available from within the 3 mm axial depth of the *µ*CT field of view. Paraffin sections were processed for H&E staining with a standard protocol^39^.

H&E-stained sections were reviewed by an expert prostate pathologist (GW) and compared with the corresponding *µ*CT sections, noting diagnostic features visible and not visible in the *µ*CT data, and any potentially valuable features observed only in the *µ*CT images.

### Alignment of *µ*CT and histology sections

For direct correlation of *µ*CT images and histology sections, a low resolution colour image of each histology section was used to search through the *µ*CT image stack using FIJI^38^ to find the best match of major structural features. In general, the orientation of the histology sections was not exactly coplanar with the transaxial *µ*CT image plane. While the isotropic *µ*CT data permits angled reslicing to enable a closer alignment of *µ*CT and histology planes, this operation introduces some interpolation artifacts. We present only native *µ*CT data in this paper in order to illustrate the inherent contrast and spatial resolution of our protocol.

### Magnetic resonance imaging

To demonstrate the potential of *µ*CT to provide 3D tissue microstructural information for validation of DWI methods, we performed 16.4 Tesla MRI microscopy on several samples prior to *µ*CT. MRI was performed at the University of Queensland node of the Australian National Imaging Facility. Briefly, prior to transfer to 80% ethanol, the gelatin-embedded tissue sample (inside the PTFE tube) was inserted into a 5-mm NMR tube (Fig.9) and equilibrated with saline containing 0.2 mL/L Magnevist gadolinium contrast agent (Bayer AG, Germany). Diffusion tensor imaging (DTI, a specific variant of DWI) was performed at spatial resolution 40*µ*m isotropic with acquisition details as described previously^33^. Fibre tractography was performed using probabilistic tensor tractography in MRtrix3 (mrtrix.org). Note that Gd contrast agent was added at low concentration to improve signal to noise ratio and not to add contrast. Any Gd likely diffused out of the samples between MRI and CT imaging sessions. We saw no evidence of Gd retention in *µ*CT or histology.

## Acknowledgments

The authors thank: Joseph Brunet, University College London, for advice on sample preparation; The facilities and scientific and technical assistance of *µ*CT beamline staff at the Australian Synchrotron; The facilities and scientific and technical assistance of the National Imaging Facility, a National Collaborative Research Infrastructure Strategy (NCRIS) capability, at the University of Queensland; The facilities and scientific and technical expertise of the Charles Perkins Centre Histology Facility at the University of Sydney.

## Author contributions statement

R.B. conceived and conducted experiments, and drafted the manuscript. B.A. and T.G. designed and optimised the *µ*CT protocol. G.W. and P.S. provided histological and surgical support. A.P. and S.D. performed histological analysis. N.K. acquired and analysed MRI data. All authors reviewed the manuscript.

